# Temperate and filamentous bacteriophages as reservoirs of bacterial virulence in stony coral tissue loss disease

**DOI:** 10.64898/2026.03.24.714041

**Authors:** Bailey A. Wallace, Lydia Baker, Erin Papke, Blake Ushijima, Stephanie M. Rosales, Cynthia B. Silveira

## Abstract

Lysogenic conversion, in which bacteriophages (viruses that infect bacteria) integrate into bacterial genomes and confer new phenotypic traits on their hosts, is a well-established mechanism for the emergence of pathogens. While this process underpins many bacterial diseases in humans and other animals, its role in coral disease remains largely unexplored. Here, we investigate the potential role of lysogenic conversion in stony coral tissue loss disease (SCTLD), which has caused unprecedented mortality of Caribbean corals since 2014 and for which the etiological agent remains unknown. Across 27 original coral samples and 178 publicly available metagenomes from the Florida Reef Tract spanning seven coral species, we detected a greater number of unique viral genomes capable of lysogeny in diseased corals or visually healthy portions of diseased corals (DD and HD, respectively) compared to visually healthy (VH) corals. SCTLD-associated bacteriophages were primarily predicted to infect bacterial taxa previously implicated in the disease, such as *Vibrionales*, *Rhodobacterales*, and *Flavobacteriia*, and carried abundant and widespread virulence genes with the potential to enhance bacterial colonization, competition, or direct host damage, including homologs of Tse2, Zot, RTX, pneumolysin, and cytolytic delta-endotoxin. Although causal relationships remain unresolved, our findings indicate that phages capable of lysogenic conversion have the genomic capacity to laterally transfer bacterial virulence genes in SCTLD-affected corals. The acquisition of genes via lysogenic conversion could contribute to bacterial virulence while maintaining community taxonomic profiles, helping to explain previous community profiling observations and providing a mechanistic framework for disease pathogenesis.

## Introduction

Since 2014, stony coral tissue loss disease (SCTLD) has devastated Caribbean and Florida reefs, affecting more than 30 species of reef-building corals and resulting in mortality rates of 40-100% in the most susceptible species (Aeby et al. 2019; Williams et al. 2021; Camacho-Vite et al. 2022). Regional patterns of SCTLD spread, characterized by clustered hotspots that propagated along the Florida Reef Tract and later throughout the Caribbean, are consistent with localized transmission from affected to nearby susceptible corals, suggesting the involvement of an infectious agent (Aeby et al. 2019; Muller et al. 2020; Traylor-Knowles et al. 2021; Camacho-Vite et al. 2022; Studivan et al. 2022). However, fine-scale patterns of spread are more nuanced, with high variability in transmission due to differences in species susceptibility and environmental conditions (Meiling et al. 2021; Williams et al. 2021; Guzmán-Urieta et al. 2026). Although some holobiont constituents have been examined as potential etiological agents, the causative agent(s) and specific mechanisms of this disease remain unknown (Landsberg et al. 2020; Ushijima et al. 2020; Veglia et al. 2022; Rosales et al. 2023; Papke et al. 2024).

Among holobiont components, the algal endosymbionts of coral (family Symbiodiniaceae) are some of the first cells to show signs of disruption upon SCTLD onset (Landsberg et al. 2020; Rossin et al. 2026). This disruption of host-Symbiodiniaceae physiology coincides with cell death and sloughing of tissues from the coral skeleton that is characteristic of SCTLD; however, it is unclear whether this disruption begins with the Symbiodiniaceae cells, coral animal cells, or other holobiont members (Landsberg et al. 2020). Viral infection has been considered as a potential mechanism underlying this disruption; however, filamentous viral-like particles observed by transmission electron microscopy are not specific to SCTLD-affected corals and instead appear to be ubiquitous across coral species and health states (Work et al. 2021; Veglia et al. 2022; Howe-Kerr et al. 2023). Another proposed mechanism of disruption is toxicosis, where free-living dinoflagellates or bacteria produce anti-eukaryotic toxins or effector molecules that could directly compromise host or symbiont cells (Miyoshi and Shinoda 2000; Landsberg 2002; Landsberg et al. 2020).

Bacterial involvement in SCTLD pathogenesis has been explored through various methods. One of the strongest pieces of evidence that bacteria may be causally involved is that lesion progression can often be slowed or arrested by the application of antibiotics (Aeby et al. 2019; Neely et al. 2020; Shilling et al. 2021; Walker et al. 2021; Forrester et al. 2024). Microbiome studies using 16S rRNA gene sequencing and metagenomics have identified enrichments of certain taxa (e.g., *Alphaproteobacteria* (*Rhodobacterales* and *Rhizobiales*), *Gammaproteobacteria* (*Vibrionales*), *Flavobacteriia* (*Flavobacterales*), and *Clostridia*) in diseased tissues, however, not all studies detect consistent microbiome shifts, and no single bacterial species has been detected universally or exclusively in disease lesions (Meyer et al. 2019; Rosales et al. 2020; Clark et al. 2021; Thome et al. 2021; Becker et al. 2022; Rosales et al. 2022; Evans et al. 2023; Rosales et al. 2023; Heinz et al. 2024; Nandi et al. 2025). Supporting a role for microbial agents in waterborne transmission, ultraviolet treatment of SCTLD-exposed water reduced the number of corals that developed lesions by 50%, which could indicate partial inactivation of bacteria or other UV-sensitive microbes (Studivan et al. 2022). Finally, cultivation-based approaches showed that bacteria isolated from SCTLD lesions displayed inconsistent ability to reproduce tissue loss in experimental assays (Ushijima et al. 2020). Therefore, none of the bacteria tested have fulfilled Koch’s postulates, by which a single isolated organism is necessary and sufficient to elicit disease in susceptible hosts, and the primary pathogen remains unknown. Together, these studies suggest that while a bacterial component of the disease is likely, additional factors that may involve complex community-level processes contribute to disease transmission and progression. These could involve dysbiotic interactions among bacterial members of the holobiont, between bacteria and the coral or Symbiodiniace cells, and between bacteria and their viruses.

Bacteriophages (phages), viruses that infect bacteria, are the most abundant symbiont of corals, yet 60 to 80% of the functions in phage genomes remain unknown (Grigson et al. 2025; Wallace et al. 2025). Phages play a critical role in shaping coral reef microbial communities through diverse infection strategies (Soffer et al. 2014; Knowles et al. 2016; Thurber et al. 2017; Breitbart et al. 2018; Wang et al. 2022; Vega Thurber et al. 2025). During lytic infection, the phage replicates viral particles and lyses its host cell, exerting a top-down control on bacterial populations, and influencing reef-wide community composition, nutrient cycling, and host-associated microbiome stability (Wilhelm and Suttle 1999; Rohwer and Thurber 2009; Barr et al. 2013; Nguyen-Kim et al. 2015; Breitbart et al. 2018; Almeida et al. 2019; Silveira et al. 2023). While lytic infection by phages has been proposed as a mechanism to actively protect coral surfaces from bacterial pathogen colonization (Barr et al. 2013; Nguyen-Kim et al. 2015; Silveira and Rohwer 2016; Almeida et al. 2019; Rubio-Portillo et al. 2024), it has also been suggested as a potential mechanism underlying SCTLD, where viral-mediated disruption of the microbial community could create ecological space for opportunistic pathogens (Nandi et al. 2025). Alternatively, temperate phages can enter a lysogenic state, integrating their genomes into the host chromosome or establishing chronic infections where their genomes persist as episomes. During lysogeny, phage genomes are replicated with the host at each cell division, and most genes involved in viral particle production are repressed (Brüssow et al. 2004). During chronic infections, which are common among filamentous bacteriophages, phage particles can be continuously produced without lysing the host cell.

Irrespective of infection type, phages can encode genes that add new or modulate existing metabolic functions of the host cell (Brüssow et al. 2004; Breitbart et al. 2007; Rohwer and Thurber 2009). Among temperate phages, these genes often mediate commensal and pathogenic interactions between bacteria and eukaryotic hosts, directly impacting host fitness and health (Brüssow et al. 2004; Duerkop et al. 2012). Notably, the role of lysogeny may be increasingly important in coral reef functioning, as reef degradation driven by anthropogenic stressors promotes microbialization, which is characterized by high microbial biomass and nutrient-altered conditions and favors lysogenic over lytic infection strategies (Haas et al. 2016; Knowles et al. 2016; Silveira and Rohwer 2016; Luque and Silveira 2020; Varona et al. 2024). Phage lysis-lysogeny decisions can therefore impact not only bacterial community structure and stability, but the overall functional capacity of the holobiont, with potential consequences for coral health, adaptability, and disease susceptibility (Cárdenas et al. 2020; Silveira et al. 2020; Voolstra et al. 2021; Voolstra et al. 2024; Baer et al. 2025; Wallace et al. 2025).

The change in genotype caused by phage genome integration into a bacterial host’s genome as a prophage (i.e. transduction) and subsequent expression of viral-encoded genes that change host phenotype is known as lysogenic conversion. Such conversion is a major mechanism for the emergence of human and animal pathogens (Wagner and Waldor 2002; Brüssow et al. 2004). This is exemplified by *Vibrio cholerae*, which has the cholera toxin genes (*ctxA*, *ctxB*) encoded by a filamentous prophage in the *Inoviridae* family (Wagner and Waldor 2002). In toxigenic *V. cholerae*, strains producing phage-encoded CTX cause severe cholera symptoms, whereas the absence of the integrated prophage may result in strains that can colonize hosts but cause only mild or asymptomatic infections (Faruque et al. 2001). Similarly, in Shiga toxin-producing *Escherichia coli* (STEC), the toxins (*stx1*, *stx2*) are encoded by a prophage belonging to *Caudoviricetes* (O’Brien et al. 1984). However, in STEC, toxin expression is tightly linked to prophage induction, meaning lysogens can exist in a non-pathogenic state until a switch from lysogenic to lytic infection occurs (Mauro and Koudelka 2011). Importantly, accumulating evidence suggests that not only can phage-bacteria interactions shape pathogenicity in facultatively free-living hosts such as *V. cholerae* and *E. coli*, but also within host-associated or intracellular niches, as in the case of *Legionella pneumophilia*, where a *Caudoviricetes* lysogen is responsible for 80% of all Legionnaire’s disease in humans (Nicholson et al. 2025). Through these mechanisms, phage-mediated virulence can be both acquired and lost, producing a spectrum of disease manifestations depending on prophage stability, induction frequency, and environmental conditions (Brüssow et al. 2004).

Bacteriophages could be involved in SCTLD through multiple mechanisms. First, a prophage-dependent pathogen may have escaped identification in previous microbiome studies, as prophage-mediated changes in pathogenicity could occur without detectable shifts in overall taxonomy and functional profiles of microbial communities. If a potential primary bacterial pathogen did exhibit phage-mediated pathogenicity, its gain and loss of prophage could cause inconsistent ability to establish infections in transmission experiments. Second, the gain and loss of bacteriophages carrying antibiotic resistance genes could contribute to the observed variability in antibiotic treatment response. Therefore, we investigate the evidence for lysogenic conversion of bacteria by bacteriophages as a potential mechanism for pathogenesis in SCTLD, where phage integration in a bacterial genome mediates the lateral transfer of genes that confer the ability of bacteria to cause disease.

## Methods

### Sample collection, processing, and sequencing

Samples were collected from three sites in the Florida Keys, USA, at a depth of approximately 7-10 m (Fig. S1) under permit number FKNMS-2022-049. These samples span three different coral species and were collected from corals with visible SCTLD tissue lesions in May of 2023 (Table S1). For each coral colony, two samples were collected: one of the tissue directly adjacent to the disease lesion (DD) and one of the visually healthy tissue from the same diseased colony (HD), at least 10 cm from the disease lesion. Upon collection, coral samples were flash-frozen and stored at −80°C until later processing. For metagenomic analysis, DNA was extracted using three different host-depletion methods: (1) a Virus and Bacteria Enrichment (VBE) method where coral and Symbiodiniaceae biomass (including DNA) are depleted through filter-fractionation and DNAse treatment, as previously described (CB samples) (Varona et al. 2023; Wallace et al. 2024), (2) whole tissue samples processed with Zymo ZR BashingBead Lysis Tubes (Zymo Research, Irvine, CA) containing 2 mm beads to lyse eukaryotic cells (ZB samples), or (3) whole tissue samples processed with Zymo ZR BashingBead Lysis Tubes with 0.1 and 0.5 mm beads to preferentially lyse bacterial cells (MB samples). Both ZB and MB samples were further processed using the HostZERO Microbial DNA Kit (Zymo Research, Irvine, CA) following the manufacturer’s instructions. The details of each extraction are outlined in the supplemental material. Paired-end (2 × 150 bp) sequencing was performed on a Novaseq platform (Illumina, San Diego, CA).

Publicly available metagenomes (N = 174) were identified through the National Center for Biotechnology Information (NCBI), using the search terms “metagenome”, “coral”, “SCTLD”, and “Stony Coral Tissue Loss” (Table S1). The inclusion of samples was limited to metagenomes of the order Scleractinia collected along the Florida Reef Tract, sequenced using Illumina, and containing associated metadata describing the geographic location of sampling, taxonomy of the coral host, and health status of the tissue sample (tissue directly adjacent to the disease lesion (DD), healthy tissue of a diseased coral (HD), and healthy tissue of a visually healthy coral (VH)). No healthy representatives of *Colpophyllia natans* from Florida were available in the public databases.

### Detection of bacteriophages with lysogenic potential

All metagenomic reads (original reads from this study and publicly available metagenomes) were adapter-trimmed and quality-filtered (k=23, trimq =30, maq = 30) using BBDuk (v39.01) (Bushnell 2014) (Fig. S2). Quality-controlled (QC) reads were assembled using MetaSpades (v3.15.5) (Nurk et al. 2017). Viral contigs, including both complete viral genomes and genome fragments (hereafter referred to as viral genomes), were identified with geNomad (v1.7.4) (Camargo et al. 2024) and VIBRANT (v1.2.1) (Kieft et al. 2020) and evaluated with CheckV (v1.0.3) (Nayfach et al. 2021). To identify phages with the potential to mediate lysogenic conversion of bacterial hosts (lysogenic conversion candidates), we screened the assembled viral genomes for signatures of temperate lifestyle (i.e., viruses that can integrate into a host genome and replicate passively without immediately lysing the host). Specifically, we focused on three categories: (1) proviruses (prophages), defined as integrated viral sequences within bacterial genomes, (2) putative temperate phages, such as those containing genes for integration, excision, and lysogen maintenance, and (3) phages known to facilitate lysogenic conversion, such as filamentous phages in the family *Inoviridae*. These groups of phages were chosen for their capacity to initiate persistent infections where they may alter host phenotypes, sometimes conferring new functions such as toxin production or enhanced host colonization (Boyd 2012). Among the broader pool of viral genomes identified by geNomad and VIBRANT (N = 179,826), five complementary approaches were applied to detect candidates: (1) geNomad’s built-in provirus prediction; (2) VIBRANT’s built-in provirus and temperate lifestyle prediction; (3) CheckV provirus classification; (4) MetaCerberus (v1.4.0) (Figueroa III et al. 2024) protein annotations (--hmm ALL), filtered using a curated list of keywords (e.g., integrase, excisionase, prophage; the complete keyword list is available via Figshare: 10.6084/m9.figshare.30853895); and (5) geNomad taxonomic identification of filamentous phages (*Inoviridae*), which were selected separately because they often lack classic hallmarks of lysogeny and can establish chronic infections while still modulating host virulence. This multi-tiered approach resulted in a final set of 6,551 phages of interest, including 407 confirmed prophages identified by geNomad, VIBRANT, or CheckV (Fig. S3a).

To assess viral presence and relative fractional abundance across samples, all phages of interest were dereplicated with MIUViG standards (95% average nucleotide identity [ANI] over 80% alignment fraction [AF]) (Roux et al. 2019) using CheckV, generating 1,384 representative viral genomes. Quality-controlled metagenomic reads were mapped back to the dereplicated viral database using Smalt (v0.7.6) at 95% identity (Wellcome Sanger). Fractional abundances of phages were then calculated using the following equation:

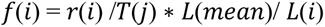

where *r*(*i*) is the number of reads mapped to each viral genome or genome fragment, *L*(*mean*) is the average genome length, *T*(*j*) represents the number of viral mapped reads in each sample, and *L*(*i*) is the length of the mapped sequence (Cobián Güemes et al. 2016). This fractional abundance calculation accounts for genome length and sequencing depth, allowing abundance comparisons in complex holobionts where different members have genome lengths with orders-of-magnitude differences (Cobián Güemes et al. 2016). A virus was considered present in a sample if it met the following thresholds: ≥ 10% genome breadth coverage and ≥ mapped reads. Presence/absence data were aggregated across samples within each disease state group (DD, HD, VH) (Fig. S3b). Importantly, a virus was considered "present" in a group even if detected in only one sample from that group. To account for how consistently phages occurred within each group, we performed an Indicator Species Analysis (ISA) (De Cáceres and Legendre 2009), which evaluates differential representation of members across groups based on both specificity and fidelity metrics (Fig. S3c; Table S2). Here, a high specificity value indicates that a phage is found primarily in a single group (i.e., DD only) or a combination of groups (i.e., DD and HD only), and a high fidelity value indicates that a phage is found across most samples in said group. All significant indicators (alpha < 0.05) were retained.

### Viral diversity

To assess the similarity between viral community profiles among the health states, the Bray-Curtis similarity algorithm was applied to viral relative fractional abundances (vegan: vegdist) and visualized with non-metric multidimensional scaling (vegan: metaMDS) (Oksanen 2013). Significant differences between groups were identified using an Adonis permutational ANOVA (adonis2; 999 permutations) stratifying by coral species and pairwise Adonis tests (Martinez Arbizu 2020). The homogeneity of multivariate dispersion was evaluated on the Bray-Curtis distance matrices (vegan: betadisper). Diversity metrics, including Shannon diversity, Simpson diversity, evenness, and richness, were also calculated among the disease states (vegan: diversity). Significant differences were detected using Kruskal-Wallis rank-sum tests for each diversity metric, adjusting p-values for multiple comparisons using the Benjamini-Hochberg false discovery rate (FDR) procedure. For indices with significant Kruskal-Wallis results, post hoc pairwise comparisons were conducted using Dunn’s tests (dunn.test) (Dinno 2024) with FDR correction applied to account for multiple testing.

### Taxonomy and functional annotations

‘Taxonomic annotation of viral genomes was derived from geNomad and functional annotation was performed with MetaCerberus using --hmm ALL (querying FOAM, KEGG, CAZy/dbCAN, VOG, pVOG, PHROG, and COG databases) as well as a custom HMM database derived from the Virulence Factor Database (VFDB; downloaded June 26, 2025). To assess differences in the relative fractional abundance of phage families and virulence gene products across disease states (DD, HD, VH), we used Kruskal-Wallis rank-sum tests and post hoc Dunn’s tests with FDR correction. Dereplicated viral genomes were combined with a dereplicated version of the NCBI Viral RefSeq database (downloaded August 8, 2025) and clustered into viral clusters (VCs) using a gene-sharing network approach with vConTACT2 (v0.9.22) (Bin Jang et al. 2019). In this framework, clusters represent viral genera, while more distant links reflect subfamily-level relationships (Bin Jang et al. 2019). The resulting network (singletons omitted) was filtered to retain only phages that met at least one of the following criteria, as well as any phages directly linked to them: (1) occurred exclusively in DD samples or in both DD and HD samples, (2) encoded at least one virulence factor (VF), or (3) were identified as disease indicator phages. The filtered network was visualized with Cytoscape (v3.10.3) (Shannon et al. 2003). This analysis provided additional taxonomic classification or confirmed *geNomad*-based assignments for those that clustered with reference phages.

### Host prediction

Phage host predictions were performed using three approaches: (1) inference from the known hosts of clustered reference phage(s), (2) iPHoP (v1.3.3) host prediction (Roux et al. 2023), and (3) for prophages, comparison of host flanking regions to the proGenomes v3 database (Fullam et al. 2023) using BLASTn (cutoffs: e-value ≤ 1e-10, alignment length ≥ 100 bp, percent identity ≥ 70%), enabling taxonomic assignments at the class level or higher (Table S3). From the resulting network, we further examined and annotated viral genomes of interest, including disease indicators, phages from families enriched in diseased samples, phages encoding virulence genes enriched in diseased samples, and phages predicted to infect bacterial hosts previously associated with SCTLD. Genomes that met these qualifications (N = 69) were annotated using SeqHub, an AI-enabled genomic context-aware platform extended from Gaia, to detect homologs of viral proteins (Jha et al. 2025). Genomes were visualized with gggenomes in R (Hackl et al. 2024).

## Results

### Viral signatures of lysogeny in SCTLD metagenomes

We identified 117 unique prophage cluster representatives in the metagenomes of the seven coral species analyzed, 112 of which were derived from diseased coral colonies (DD or HD). While the presence of flanking host sequences can confirm prophage status, many true proviruses may lack sufficiently covered flanking regions in metagenomic data for assembly and detection. Therefore, we examined an additional 1,640 viruses as they encoded genes associated with a temperate lifestyle (enabling lysogenic infections) or if they encoded structural genes from viral taxonomic groups commonly associated with lysogenic conversion of bacterial hosts, such as the family *Inoviridae*. These included genes related to phage integration (e.g., integrases, excisionases), virulence (e.g., toxins, effectors, host colonization), filamentous virus structure (e.g., minor coat proteins, ssDNA binding proteins, and secretin/extrusion channel proteins), and lysogen maintenance and stability (e.g., toxin-antitoxin systems). Among all viruses with lysogenic potential (including prophages, temperate viruses, and *Inoviridae*; herein referred to as lysogenic conversion candidates), 137 were exclusive to DD samples, and 99 were identified in both DD and HD samples (Fig. S3b). Only seven lysogenic conversion candidates were exclusive to VH samples, and most viruses (N = 1,462) were present in all three groups.

The ISA identified a total of nine indicator viruses associated with diseased samples (Fig. S3c). Potential pathogenic viruses were expected to have high specificity and fidelity in DD samples or in both DD and HD samples (where disease has emerged or is partially established), but not in HD samples alone. Two viruses were unique indicators of disease lesion (DD) samples, and seven viruses were shared between DD and HD samples. The most prominent indicator virus, SINT_DD_R2S43C_NODE_111 (class *Caudoviricetes*), exhibited 96% specificity to the DD group and 64% fidelity, meaning it was almost exclusively found in DD samples (specificity) and present in the majority of DD samples (fidelity), despite inherent challenges in viral detection. No indicator viruses were identified for HD samples.

### Community profiles of lysogenic conversion candidates

The communities of lysogenic conversion candidates, i.e., viruses with lysogenic potential, including prophages, temperate viruses, and *Inoviridae*, were significantly different between the three disease states (DD, HD, and VH; PERMANOVA, R² = 0.041, pseudo-F = 4.25, p = 0.033), after controlling for species-level differences (Fig. 1a). When species was included as a fixed factor, the effect of disease state remained significant (R² = 0.013, pseudo-F = 2.21, p = 0.019), reinforcing the influence of disease state on lysogenic conversion candidate communities. While these results also indicated that species was a significant driver of group differences (R² = 0.401, pseudo-F = 23.04, p = 0.001), it is important to note that the number of replicates per species varied across disease state categories (Table S4), and beta dispersion among species groups was significantly different (PERMDISP, F = 2.98, df = 6, p = 0.008). As a result, uneven replication among species and differences in dispersion may influence these patterns. Pairwise comparisons revealed that HD samples differed significantly from both DD (R² = 0.042, pseudo-F = 6.49, p = 0.001) and VH (R² = 0.053, pseudo-F = 4.61, p = 0.001) samples, while DD and VH were not significantly different (R² = 0.012, pseudo-F = 1.98, p = 0.057) (Fig. 1a).

**Fig. 1.**
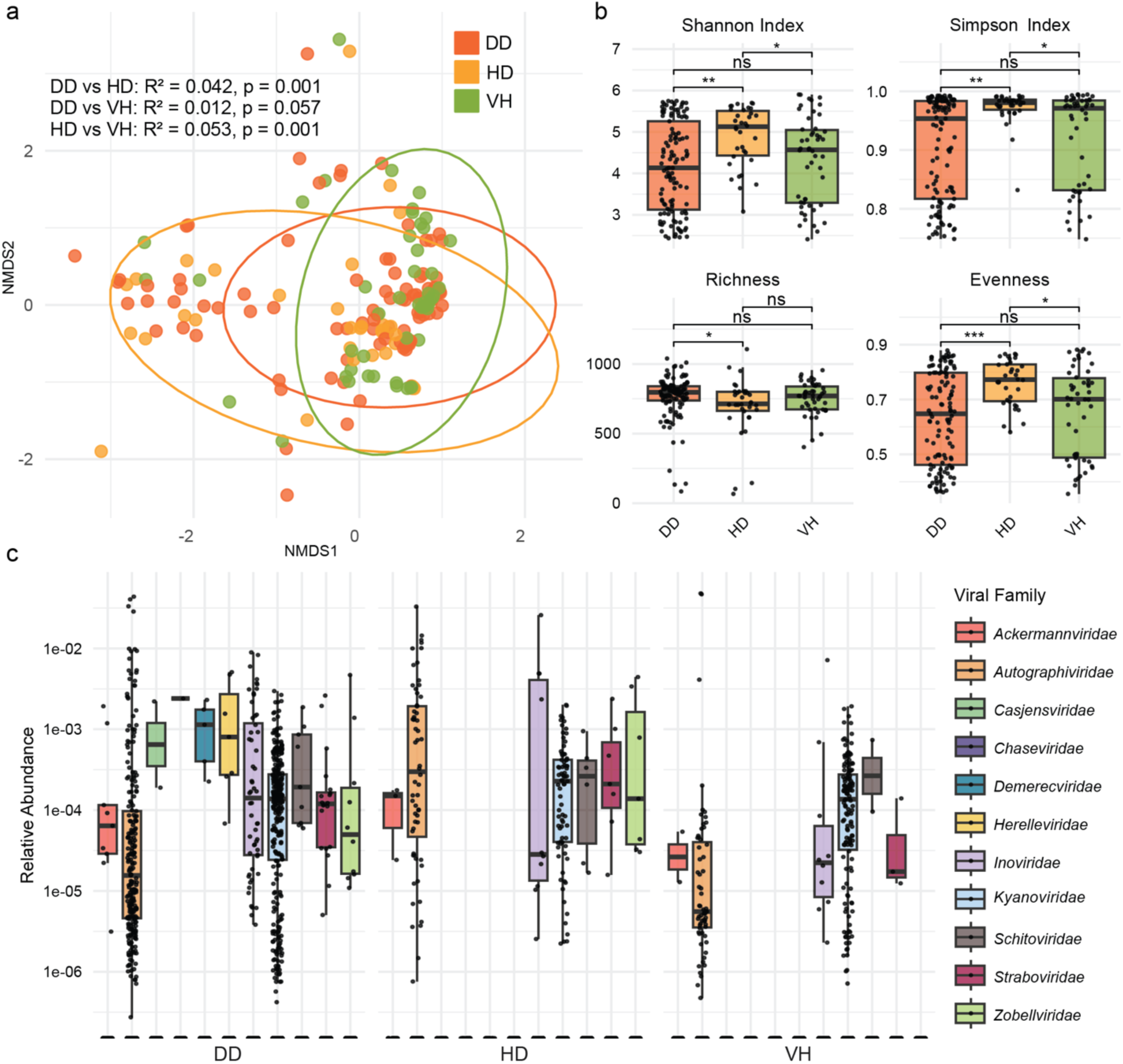
Lysogenic conversion candidate community differences between SCLTD-infected lesion-adjacent tissue (DD), healthy tissue of diseased corals (HD), and healthy tissue of healthy corals (VH). (a) Non-metric multidimensional scaling (NMDS) plot by bacterial genus. The NMDS plot is based on Bray-Curtis distances calculated from the relative fractional abundances of individual viruses and utilizes 999 permutations. Circles denote a 95% CI for the clustered points. (b) Mean alpha-diversity indices of lysogenic conversion candidate virus communities for each of the three groups, including Shannon Index, Simpson Index, Richness, and Evenness. Significance codes: 0.001 ‘**’, 0.01 ‘*’, and not significant “ns”. (c) Relative fractional abundances of temperate virus families by disease state

The alpha-diversity metrics displayed similar patterns, where HD samples were distinct from DD and VH samples (Fig. 1b). The mean Shannon Index of HD samples (4.89 ± 0.12) was significantly higher than DD samples (4.15 ± 0.10; p.adj = 0.001) and VH samples (4.33 ± 0.15; p.adj = 0.035). Mean Simpson Index was also elevated in HD samples (0.97 ± 0.005), relative to DD (0.91 ± 0.008; p.adj = 0.002) and VH (0.92 ± 0.012; p.adj = 0.032). Mean viral genome richness was significantly lower in HD samples (690.71 ± 37.65) than DD samples (762.12 ± 14.63; p.adj = 0.023) but was similar to VH samples (752.32 ± 16.31). Viral genome evenness was also elevated in the HD samples (0.76 ± 0.01) compared to DD samples (0.63 ± 0.02; p.adj < 0.001) and VH samples (0.66 ± 0.02; p.adj = 0.012).

A taxonomic analysis of all lysogenic conversion candidates showed interesting patterns at the family level (Fig. 1c). More bacteriophage families were identified in the DD samples (N = 11) than in HD (N = 7) or VH (N = 6). A Kruskal-Wallis test with FDR correction indicated significant differences in relative abundance of *Autographiviridae* (χ² = 13.3, df = 2, p = 0.005), *Zobellviridae* (χ² = 16.8, df = 2, p = 0.001), and *Inoviridae* (χ² = 8.3, df = 2, p = 0.046) across disease states (Table S5). A post-hoc pairwise Dunn’s test with FDR correction showed that DD samples had higher *Autographiviridae* abundances (belonging to the tailed phage group) compared to VH (DD vs VH: p.adj < 0.001), while no significant difference was observed between DD and HD samples (p.adj = 0.461), or VH and HD (p.adj = 0.051). *Autographiviridae* were also more prevalent in DD samples (occurring in 77.78% of the samples) compared to HD (41.18%) and VH (56.00%). *Zobellviridae*, another family in the class *Caudoviricetes* (tailed phages), was only detectable in DD and HD samples. This family was significantly more abundant in HD samples than in DD samples (p.adj = 0.009) and VH samples (p.adj < 0.001) and differed between DD and VH samples (p.adj = 0.023). For *Inoviridae*, pairwise tests revealed significantly higher abundance in DD samples compared to VH samples (p.adj = 0.018), but no differences were detected between DD and HD (p.adj = 0.174) or HD and VH (p.adj = 0.477). *Inoviridae* prevalence was also higher in the DD samples (34.19%) compared to HD (17.65%) and VH (18.00%). The *Caudoviricetes* families *Casjensviridae*, *Chaseviridae*, *Demerecviridae*, and *Herelleviridae* were detectable in DD samples only, but were found in very few samples (1.71%, 0.85%, 4.27%, and 3.42% of DD samples, respectively). In contrast, *Kyanoviridae*, another tailed bacteriophage, showed similar abundance (χ² = 1.4, df = 2, p = 0.589) and prevalence (DD: 94.02%, HD: 85.29%, and VH: 96.00%), regardless of disease state.

### Virulence genes in lysogenic conversion candidate viruses

The diversity of genes with homology to virulence genes in known pathogens encoded by lysogenic conversion candidates was greater in DD samples (N = 40) compared to HD (N = 38) and VH (N = 34) (Fig. 2). Several of these genes were found exclusively in the SCTLD-infected coral samples (DD and/or HD), including genes encoding for putative virulence factor homologs, such as a Zonula occludens toxin (Zot) like proteins, iron-regulated protein FrpC (RTX protein family), cholesterol-dependent cytolysin lly (Pneumolysin-like), and virulence factors related to bacterial lifestyle regulation and adaptation, such as poly-beta-1,6 N-acetyl-D-glucosamine synthase PgaC (PNAG), and carbon storage regulator A (CsrA). However, these exclusively SCTLD-associated virulence genes were not always highly prevalent, as they were detectable in only 0.85% to 6.84% of DD samples and 2.94% to 5.88% of HD samples (Table S6).

**Fig. 2.**
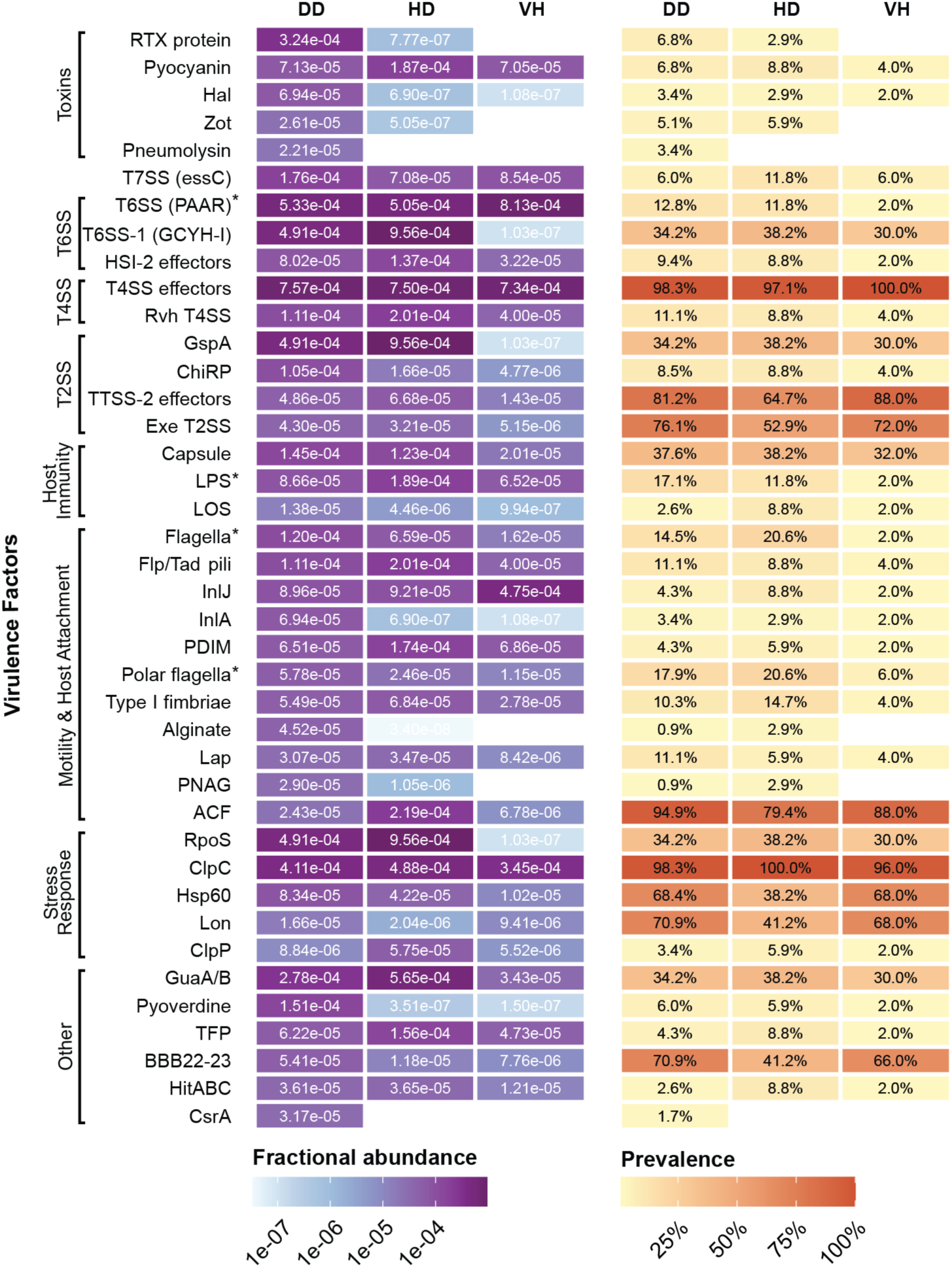
Lysogenic conversion candidate-encoded virulence genes. Fractional abundance and prevalence of virulence genes encoded by lysogenic conversion candidate viruses are shown across disease states (DD, HD, and VH). Virulence genes were identified using a hidden Markov model (HMM) database derived from the Virulence Factor Database (VFDB). Asterisks (*) indicate significant differences in fractional abundances between disease states

Six of the other 35 identified genes with homology to virulence genes in known pathogens had significant differences in relative abundance across disease states. Genes annotated as “LPS” in VFDB, representing functions related to (but not exclusive of) lipopolysaccharide-mediated virulence (e.g., immune evasion, complement resistance, or modulation of host responses), were enriched in both HD (1.89e-04 ± 8.98e-05; mean, SE) and DD (8.66e-05 ± 1.83e-05) samples, compared to VH (6.52e-05 ± 2.96e-05; both p.adj < 0.001). Motility-related genes showed a similar pattern: polar flagella were more abundant in HD (2.46e-05 ± 8.58e-06) and DD (5.78e-05 ± 1.85e-05) than in VH (1.15e-05 ± 5.34e-06; both p.adj < 0.001), and flagella was higher in HD (6.59e-05 ± 3.98e- 05) and DD (1.20e-04 ± 5.92e-05) than VH (1.62e-05 ± 1.58e-05; p.adj = 0.003 and 0.016, respectively). In contrast, Type VI secretion system (T6SS) genes were significantly reduced in HD (5.05e-04 ± 2.29e-04) and DD (5.33e-04 ± 1.72e-04) relative to VH (8.13e-04 ± 4.73e-04; p.adj = 0.012 and 0.006, respectively). Type I fimbriae were elevated in HD compared to both VH (6.84e-05 ± 2.60e-05 vs. 2.78e-05 ± 1.38e-05; p.adj = 0.014) and DD (5.49e-05 ± 3.00e-05; p.adj = 0.040). Finally, Hsp60 was more abundant in HD than VH (4.22e-05 ± 2.35e-05 vs. 1.02e-05 ± 6.95e-06; p.adj = 0.027), but lower in HD than DD (8.34e-05 ± 3.81e-05; p.adj = 0.047). We also examined taxonomic and virulence gene patterns split by coral species, but due to variation in the number of replicates per species within each disease state, all samples were grouped for analysis (Fig. S4; Fig. S5).

### Genome similarity network of lysogenic conversion candidates

The pool of lysogenic conversion candidates (N = 1,757) was reduced to include only the viruses that were potential agents of disease. This reduced group of viruses included (1) viruses exclusively found in SCTLD-infected corals (DD or HD), (2) viruses encoding virulence factors, or (3) viruses identified disease indicators, and are herein referred to as SCTLD-associated lysogenic conversion candidates. These viruses were visualized in a genome-wide gene-sharing network, along with their nearest neighbors (N = 231 total viruses), and were represented in at least nine different bacteriophage families (Fig. 3a; Table S3). The majority of SCTLD-associated lysogenic conversion candidates were only classified at the class level (dsDNA-tailed phages; *Caudoviricetes*), while a subset could be identified at the family level, including *Autographiviridae*, *Herelleviridae*, *Steigviridae*, *Winoviridae*, *Zobellviridae*, *Mushuvirus*, *Muvirus*, *Schmittlotzvirus*, and *Inoviridae*). Inferred bacterial hosts of these SCTLD-associated viruses included members of the classes *Alphaproteobacteria*, *Clostridia*, *Bacteroidia*, *Gammaproteobacteria*, *Desulfovibrionia*, *Betaproteobacteria*, *Bacillota*, *Winogradskyella*, *Oligoflexia*, and *Deinococci*. Three of the nine disease indicator viruses were present in this network: SINT_DD_R2S4C3D_NODE_111 clustered with a *Vibrio*-infecting *Caudoviricetes* phage, OFAV_DD_S-3342_NODE_52649 clustered with a group of unknown viruses, and SINT_DD_R2S4C3D_NODE_62 was unclustered, but more distantly linked to a group of *Clostridia*-infecting *Caudoviricetes* phages.

**Fig. 3.**
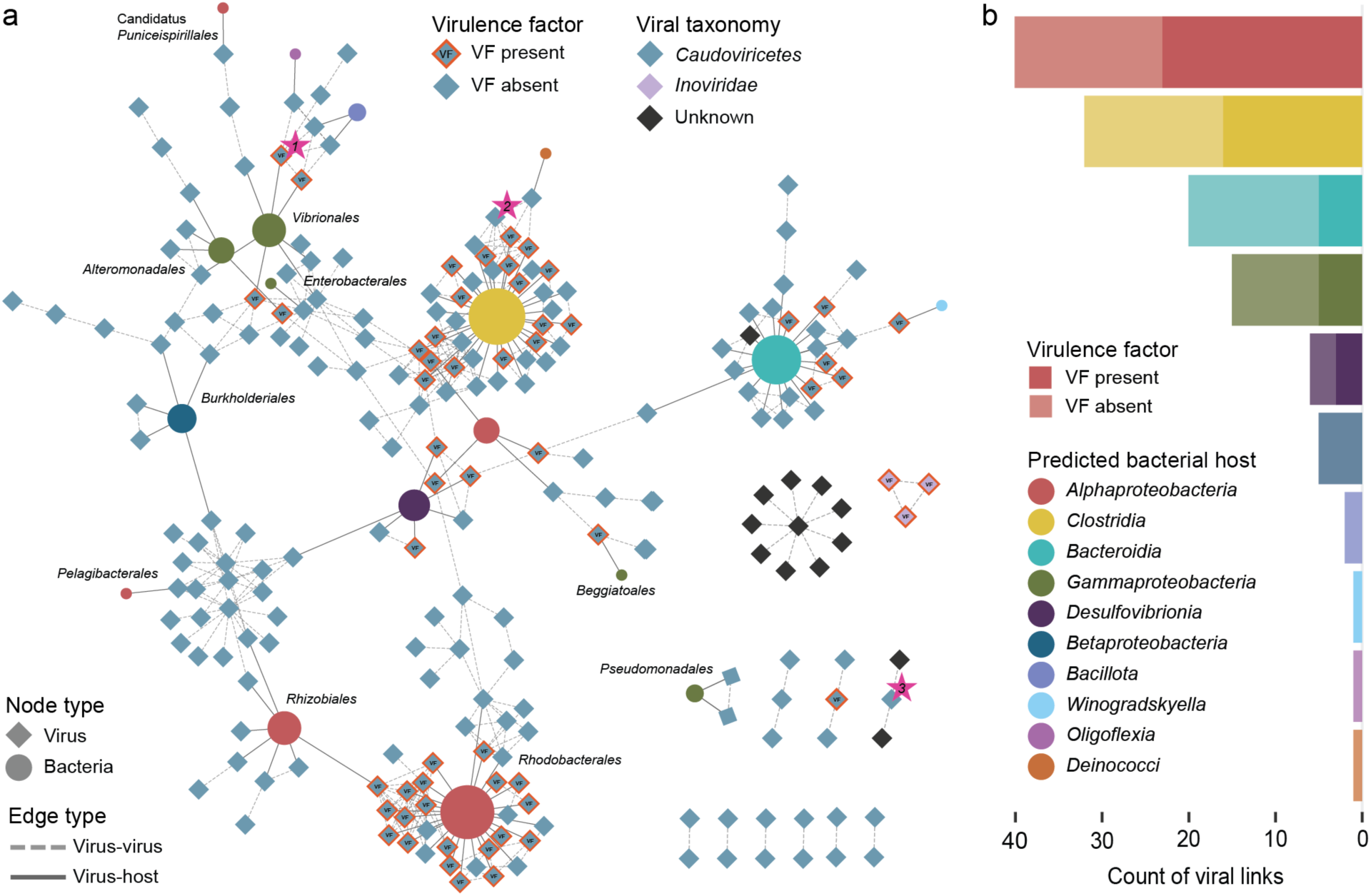
Gene sharing network and host associations of SCTLD-associated lysogenic conversion candidates. (a) Lysogenic conversion candidates associated with SCTLD-infected corals were clustered into a gene-sharing network at the genus level. Each node represents an individual virus (diamond) or a predicted bacterial host (circle), and host nodes are sized based on the number of viral linkages. Dashed edges indicate links between viruses and solid edges indicate virus-host links. Node fill colors indicate taxonomy (class/family level for viral nodes and class level for bacterial nodes). Some bacterial host assignments were distinguished at the order level (indicated by text labels), and those with no label were grouped at the class level. Orange outlines and black “VF” labels on viral nodes denote the presence of at least virulence factor (VF) within a given viral genome and disease indicator viruses in the network are shown with small pink stars labeled (1) SINT_DD_R2S4C3D_NODE_111, (2) SINT_DD_R2S4C3D_NODE_62, and (3) OFAV_DD_S-3342_NODE_52649. (b) Count of viral links by bacterial host class. Stacked bar chart shows the total number of viral links identified for each bacterial class (opaque segments) versus the counts of links where the virus encoded a predicted virulence factor (VF; semi-transparent segments). Bacterial classes are ordered from lowest to highest total count

Three viruses in the family *Inoviridae* were present in the network, all of which were present only in SCTLD-infected samples and contained putative virulence genes. Several predicted host taxa previously associated with SCTLD (*Vibrionales* (*Gammaproteobacteria*), *Rhodobacterales* (*Alphaproteobacteria*), *Flavobacteriia* (*Bacteroidia*), and *Clostridia* (Meyer et al. 2019; Rosales et al. 2020; Clark et al. 2021; Becker et al. 2022; Rosales et al. 2022; Rosales et al. 2023) were identified here and were linked to virulence gene-encoding temperate phages or prophages. Of the identified host classes, *Alphaproteobacteria* was most frequently linked to SCTLD-associated lysogenic conversion candidates (N = 40), followed by *Clostridia* (N = 32), and *Bacteroidia* (N = 20) (Fig. 3b). *Alphaproteobacteria* exhibited the highest proportion of VF-encoding links (58% of its associated viruses encoded VFs), followed by *Clostridia* (50%) and *Desulfovibrionia* (50%), which had comparatively fewer total links (N = 6).

### Virulence gene-encoding bacteriophages associated with SCTLD

To further narrow the search for viruses capable of modulating pathogenesis in SCTLD, we focused on a subset of the SCTLD-associated lysogenic conversion candidates that were (1) linked to bacterial hosts previously implicated in the disease, (2) members of viral families enriched in SCTLD-infected corals, or (3) encoding putative virulence genes enriched in SCTLD-infected corals. Among the genomes of these SCTLD-pathogenicity candidates (N = 65), multiple virulence factor functional categories were represented, including toxins, secretion system components, host colonization factors, and persistence genes related to survival within host cells (Table S7). A subset of SCTLD-pathogenicity candidates (N = 13 individual viruses) carried virulence factors in the “Toxins/Effectors & Effector Delivery” category (Table 1), which were of particular interest due to their potential to directly damage host cells (coral or endosymbiotic algae). Notably, several of these viruses encoded genes with homology to known bacterial toxins, including Toxin Tse2 (*tse2*), Exotoxin type G (*speG*), Type-2Bb cytolytic delta-endotoxin (*cyt2Bb1*), Heat-stable enterotoxin A (*ystA*), Peptidoglycan amidase Tse1 (*tse1*), Zonula occludens toxin (*zot*), Accessory cholera enterotoxin (*ace*), Iron-regulated RTX toxin (*frpA*), and a thiol-activated cytolysin annotated as a Lectinolysin (*lly*) or Seeligeriolysin (*lso*) homolog. In addition, viruses carried genes for secretion systems and effectors, including components of the type II secretion system (*exeD*), type VII secretion system (*essC*, *essA*), as well as type IVB Dot/Icm system (*coxFIC1*).

**Table 1.** Viral candidates for lysogenic conversion in SCTLD pathogenesis. Candidates for lysogenic conversion include disease indicator viruses, viruses that infect SCLTD-associated bacterial hosts, and viruses that are associated with viral families or virulence factors that were enriched in SCTLD-infected samples. The ID, cluster, taxonomy, type, virulence genes, virulence categories, and host information are reported for these viruses

| Virus ID | Cluster | Taxonomy | Type | Virulence gene(s) | Category | Host |
| --- | --- | --- | --- | --- | --- | --- |
| SINT_DD_R2S4C3D_NODE_111* | VC_170_0 | *Caudoviricetes* | Temperate | (tse2) Toxin Tse2 | Toxins/Effectors & Effector Delivery | Vibrionales |
| MCAV_VH_McLGI_NODE_221 | VC_190_0 | *Mushuvirus (Caudoviricetes)* | Temperate | (essC) type VII secretion system protein<br>(sap) S-layer protein<br>(ide) IgM protease | Toxins/Effectors & Effector Delivery<br>Survival in Host & Immune Modulation<br>Survival in Host & Immune Modulation | *Clostridia* |
| MCAV_HD_McBI_NODE_339 | VC_52_0 | *Winoviridae (Caudoviricetes)* | Provirus | ESAT-6 secretion machinery protein EssA | Toxins/Effectors & Effector Delivery | *Flavobacteriia* |
| MCAV_DD_McD6_NODE_117 | VC_52_0 | *Winoviridae (Caudoviricetes)* | Provirus | ESAT-6 secretion machinery protein EssA<br>Depolymerase, capsule K11-specific<br>Exotoxin type G<br>Host translation inhibitor 5b<br>Type-2Bb cytolytic delta-endotoxin<br>Heat-stable enterotoxin A | Toxins/Effectors & Effector Delivery<br>Motility & Host Colonization<br>Toxins/Effectors & Effector Delivery<br>Survival in Host & Immune Modulation<br>Toxins/Effectors & Effector Delivery<br>Toxins/Effectors & Effector Delivery | *Flavobacteriia* |
| SINT_DD_R2S4C3D_NODE_272 | VC_236_0 | *Caudoviricetes* | Temperate | (coxFIC1) Coxiella Dot/Icm type IVB secretion system translocated effector<br>(FlhD) Flagellar transcriptional regulator | Toxins/Effectors & Effector Delivery<br>Motility & Host Colonization | *Rhodobacteriales* |
| SINT_DD_R2S4C3D_NODE_12 | VC_241_0 | *Caudoviricetes* | Provirus | (Tse1) Peptidoglycan amidase<br>(FlhD) Flagellar transcriptional regulator<br>(acpXL) acyl carrier protein<br>(flmH) short chain dehydrogenase/reductase family oxidoreductase | Toxins/Effectors & Effector Delivery<br>Motility/Host Colonization<br>Survival in Host & Immune Modulation<br>Motility & Host Colonization | *Rhodobacteriales* |
| SINT_DD_R2S4C1D_NODE_3979 | VC_1483_0 | *Inoviridae (Faserviricetes)* | Temperate/Other | (exeD) general secretion pathway protein D<br>(zot) Zona occludens toxin | Toxins/Effectors & Effector Delivery<br>Toxins/Effectors & Effector Delivery | *Unknown* |
| SINT_DD_R2S4C2D_NODE_1517 | VC_1483_0 | *Inoviridae (Faserviricetes)* | Provirus | (exeD) general secretion pathway protein D<br>(zot) zona occludens toxin | Toxins/Effectors & Effector Delivery<br>Toxins/Effectors & Effector Delivery | *Unknown* |
| SINT_DD_R2S4C2D_NODE_577 | VC_1483_0 | *Inoviridae (Faserviricetes)* | Temperate/Other | (exeD) general secretion pathway protein D<br>(zot) Zona occludens toxin | Toxins/Effectors & Effector Delivery<br>Toxins/Effectors & Effector Delivery | *Unknown* |
| MBOFD34DD_S3_NODE_16537 | Singleton | *Inoviridae (Faserviricetes)* | Temperate/Other | (zot) zona occludens toxin<br>(ace) Accessory cholera enterotoxin | Toxins/Effectors & Effector Delivery<br>Toxins/Effectors & Effector Delivery | *Unknown* |
| SINT_DD_R2S4C4D_NODE_758 | Singleton | *Unknown* | Temperate | (zot) zona occludens toxin<br>(exeD) general secretion pathway protein D<br>(ace) Accessory cholera enterotoxin | Toxins/Effectors & Effector Delivery<br>Toxins/Effectors & Effector Delivery<br>Toxins/Effectors & Effector Delivery | *Unknown* |
| SINT_DD_R2S4C2D_NODE_1892 | Singleton | *Caudoviricetes* | Temperate | (frpA) iron-regulated protein FrpC [RTX protein] | Toxins/Effectors & Effector Delivery | *Unknown* |
| DLAB_DD_R2S4B3D_NODE_5674 | Singleton | *Caudoviricetes* | Temperate | (lly) lectinolysin [Pneumolysin]; (lso) Seeligeriolysin | Toxins/Effectors & Effector Delivery | *Unknown* |

Genome plots of nine viruses encoding virulence factors with potential to damage eukaryotic cells revealed diverse host associations and virulence functions (Fig. 4a). A *Caudoviricetes* virus (1. SINT_DD_R2S43C_NODE_111) infecting *Vibrio* and identified as an indicator virus of SCTLD (96% specificity and 64% fidelity to the DD group), encoded a gene with homology to the bacterial virulence toxin Tse2. Another *Caudoviricetes* (2. MCAV_DD_McD6_NODE_117) was identified as a prophage likely infecting *Flavobacteriia*. This virus encoded genes with homology to exotoxin type G, type-2Bb cytolytic delta-endotoxin, and heat-stable enterotoxin A. Four filamentous bacteriophages in the family *Inoviridae* were identified here (3. SINT_DD_R2S4C1D_NODE_3979, 4. SINT_DD_R2S4C2D_NODE_1517, 5. SINT_DD_R2S4C2D_NODE_557, and 6. MBOFD34DD_S3_NODE_16537). These viruses all encoded homologs to zonula occludens toxin, three of the four encoded homologs to general secretion pathway protein D, and one encoded a homolog to accessory cholera enterotoxin. A single virus (7. SINT_DD_R2S4C4D_NODE_758) encoded all three of these virulence genes (zona occludens toxin, general secretion pathway protein D, and accessory cholera enterotoxin) but had no identified host. Two *Caudoviricetes* viruses with no identified hosts (SINT_DD_R2S4C2D_NODE_1892 and DLAB_DD_R2S4B3D_NODE_5674) encoded homologs to iron-regulated RTX toxin and a pneumolysin/seeligeriolysin, respectively.

**Fig. 4.**
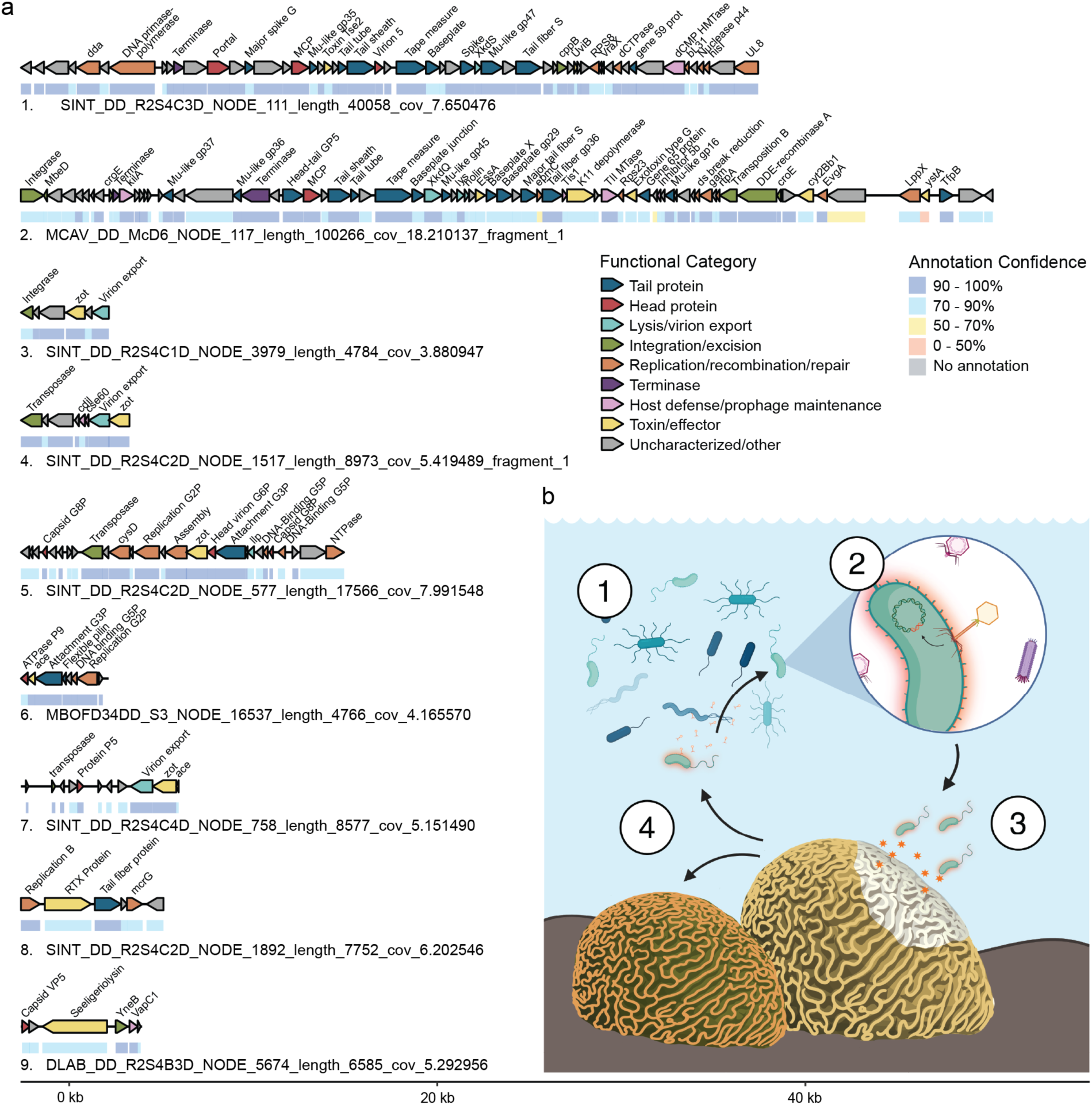
Genomes of toxin-encoding lysogenic conversion candidate viruses and conceptual hypothesis of lysogenic conversion in SCTLD pathogenesis. (a) Genome maps of temperate viruses encoding putative toxin genes. Annotations highlight genes related to phage structure, lysogeny, replication, and toxins. (b) Conceptual model illustrating the potential role of lysogenic conversion in the progression of stony coral tissue loss disease (SCTLD). (1) Bacteria in the surrounding environment interact with phages released from other infected bacteria. (2) The phage integrates its genome into the bacterial host chromosome. (3) Through lysogenic conversion, the host bacterium expresses virus-encoded virulence factors or toxins, enhancing colonization of the coral holobiont and/or directly harming coral or Symbiodiniaceae cells. (4) Infected bacteria and/or viruses are then dispersed via water, sediments, or direct contact, facilitating further infection and disease transmission. This conceptual figure incorporates elements created with BioRender.com

## Discussion

Identifying the causative agents of coral diseases such as SCTLD remains a major challenge, in part due to the complexity of the coral holobiont. Here, we explore the possibility that bacteriophages play a role in SCTLD etiology via lysogenic conversion, whether through the emergence of a primary pathogen or by facilitation of opportunistic microbial growth. Phages are integral to bacterial ecology, modulating host abundance, gene content, and virulence in numerous systems and diseases (Wagner and Waldor 2002; Boyd 2012). Lysogenic conversion, in which temperate phages integrate into bacterial genomes and alter host traits, has previously been proposed as a plausible mechanism for the emergence of pathogenic bacterial phenotypes in coral reefs (Weynberg et al. 2015). Metagenomics provides a unique opportunity to investigate viral contributions, expanding on what we have learned through culture-based surveys and marker gene studies. In the present study, we show the genomic contributions of temperate bacteriophages in SCTLD-affected corals, shedding light on an overlooked layer of complexity that may contribute to the difficulty in identifying consistent bacterial pathogens associated with this disease.

### Temperate viruses are abundant in SCTLD-affected coral microbiomes

Viral signatures of lysogeny were detected across seven coral species affected by SCTLD. Though the majority of temperate viruses (85%) were present in the metagenomes of both healthy and diseased corals, nearly 15% were exclusive to the diseased coral samples (DD or HD; Fig. S3b), and only 0.4% were exclusive to healthy corals (VH), suggesting that temperate phages are enriched in disease states. Given that lysogeny is a common viral strategy under host-rich conditions (Knowles et al. 2016), its higher prevalence here may reflect a shift in bacteria-phage dynamics driven by disease-associated microbial proliferation. Although changes in physical microbial density have not yet been quantified as SCTLD lesions progress, it is likely that the release of nutrients and cellular debris during tissue loss promotes the proliferation of opportunistic heterotrophs, increases bacterial densities, and encourages lysogenic infection (Barott and Rohwer 2012; Haas et al. 2016; Knowles et al. 2016; Silveira and Rohwer 2016). Importantly, increased bacterial loads in this context would not necessarily indicate causation; they may instead represent opportunistic secondary infections following the physiological breakdown, which then help sustain the disease microenvironment.

### Inoviridae and Autographiviridae are enriched in SCTLD-infected corals

Temperate viral communities differed across the disease states at the individual virus level (Fig. 1a), with HD samples showing distinct composition relative to DD and VH corals. HD communities were more even and diverse, which may represent a restructuring of viral assemblages in the visibly healthy tissues of diseased colonies (Fig. 1b). This may mimic what has been observed in microbial communities, where disease-unaffected (DU; here equivalent to HD) samples had the highest alpha-diversity and a distinct microbial composition (Rosales et al. 2023), suggesting that SCTLD may disrupt the microbiome prior to gross lesion formation. Importantly, the increase in HD alpha-diversity could also reflect variability in sample collection distance from lesions (Rosales et al. 2023). In DD samples, viral species richness was higher than in HD samples (Fig. 1b), a trend that was also reflected at the family-level taxonomy.

The viral families *Autographiviridae* and *Inoviridae* were more abundant and prevalent in disease-associated samples (Fig. 1c), *Zobellviridae* was enriched in HD samples, and several *Caudoviricetes* families were detectable only in DD samples, albeit at low prevalence. Most members of *Autographiviridae,* dsDNA tailed phages in *Caudoviricetes*, are strictly lytic (Molineux 2006; Boeckman et al. 2022), though recent work has shown that T7-like *Autographiviridae* phages carrying lysogeny modules also exist (Boeckman et al. 2022; Putzeys et al. 2023).

These viruses infect hosts such as *Acinetobacter baumannii*, *Roseobacter* strains, *Pseudomonas*, *Desulfovibrio*, and *Aeromonas* (Boeckman et al. 2022; Bujak et al. 2022; Putzeys et al. 2023; Du et al. 2025; Liu et al. 2025). Strains of *Autographiviridae* have been investigated for their potential therapeutic use against multidrug-resistant *Acinetobacter baumannii*, a pathogen causing skin and soft tissue infections such as pyoderma and necrotizing fasciitis (Wintachai et al. 2022; Liu et al. 2025). While these bacterial hosts are not necessarily present in corals, their necrotizing capabilities illustrate that such phages can interact with bacteria capable of tissue damage.

*Inoviridae* are common agents of lysogenic conversion, establishing chronic infections where they release viral progeny over time without killing the host cell. They are known to primarily infect Gram-negative bacteria, including species that are pathogens of humans (*Vibrio cholerae*, *Pseudomonas aeruginosa*, *Neisseria meningitidis*), plants (*Ralstonia solanacearum*), and animals (*Vibrio anguillarum*), illustrating their capacity to modulate bacterial virulence (Waldor and Mekalanos 1996; Yamada 2013; Bille et al. 2017; Burgener et al. 2019; Mauritzen et al. 2020; Weng et al. 2025). Chronic infection by *Inoviridae* viruses enables them to subtly modulate host physiology, including promoting biofilm formation, altering motility, and enhancing virulence traits, which can fundamentally reshape microbial community dynamics (Waldor and Mekalanos 1996; Bille et al. 2017; Burgener et al. 2019). Here, *Inoviridae* were more prevalent and abundant in DD and HD samples compared to VH, suggesting that they may contribute to bacterial community virulence in diseased corals. These findings highlight that SCTLD influences both fine-scale (individual virus) and broader (family-level) structures of temperate viral assemblages, likely mirroring shifts in host bacterial populations.

### A genome similarity network reveals viruses infecting key bacterial symbionts

A protein sharing network analysis of 231 SCTLD-associated lysogenic conversion candidate viruses (including virulence gene-encoding viruses and disease indicators) spanned at least nine bacteriophage families (Fig. 3a). Most viruses were classified at the class level (*Caudoviricetes*), and the majority of predicted bacterial hosts included taxa previously linked to SCTLD, including *Vibrionales* (*Gammaproteobacteria*), *Rhodobacterales* (*Alphaproteobacteria*), *Flavobacteriia*, *Clostridia*, and *Bacteroidia* (Meyer et al. 2019; Rosales et al. 2020; Clark et al. 2021; Becker et al. 2022; Rosales et al. 2022; Rosales et al. 2023). Notably, *Inoviridae* viruses in this disease-associated network carried multiple virulence genes, highlighting a potential mechanism for targeted modulation of disease-associated bacteria.

Among hosts in the network, *Alphaproteobacteria* and *Clostridia* stood out, with both a high absolute number of linkages to SCTLD-associated viruses (N = 40 and N = 32, respectively), and a high proportion of VF-encoding viruses (57% and 50%, respectively), suggesting that members of these classes may be disproportionately affected by lysogenic conversion events (Fig. 3b). Within the *Alphaproteobacteria*, this signal was driven primarily by *Rhodobacterales*, a lineage commonly abundant in coral microbiomes and frequently associated with disease states (Sunagawa et al. 2009; Clark et al. 2021; Rosales et al. 2023). As members of *Alphaproteobacteria* and *Clostridia* are frequently infected by temperate viruses that carry virulence-associated genes in this dataset, viral integration could meaningfully alter their functional potential within the coral microbiome. Given that shifts in *Alphaproteobacteria* and *Clostridia* have been observed in other coral disease contexts (Sunagawa et al. 2009; Mhuantong et al. 2019) as well as in SCTLD (Meyer et al. 2019; Clark et al. 2021; Rosales et al. 2023), their strong association with VF-encoding, SCTLD-associated temperate viruses raises the possibility that phage-mediated gene transfer may contribute to pathogenic transitions or community destabilization.

### Evidence for phage-mediated bacterial virulence in SCTLD

Potential virulence factors (VFs) were more abundant, prevalent, and diverse in DD and HD samples compared to VH. Among the lysogenic conversion candidates (proviruses, temperate viruses, and filamentous viruses) in SCTLD-infected and healthy corals, 40 unique virulence factors were detected (Fig. 2). These phage-encoded genes were functionally diverse: some promote bacterial colonization of hosts (e.g., fimbriae, pili, capsules, flagella), others enhance bacterial competition and survival (e.g., T6SS, T4SS, stress response genes), and a subset could more directly increase bacterial virulence (e.g., Zot, Pneumolysin, RTX, Pyocyanin homologs). Several VFs were detected exclusively in DD samples, albeit at low prevalence, which may reflect either detection limits or genuine heterogeneity among colonies. This included several homologs to toxins such as Zot, Pneumolysin, and RTX protein, which could potentially damage eukaryotic coral cells directly (Fasano et al. 1995; Cockeran et al. 2002; Frey and Kuhnert 2002). Enrichment of VFs in diseased corals may reflect proliferation of opportunistic or primary pathogens, some leveraging phage-encoded functions for attachment, motility, or pathogenicity. Even at low prevalence, the presence of these virulence genes uniquely in SCTLD-infected corals highlights that viral genomes may contribute to disease-associated microbial dynamics and potentially lesion formation.

Thirteen SCTLD-associated lysogenic conversion candidates encoded VFs that may directly damage or disrupt the physiology of eukaryotic cells, including homologs of Tse2, cytolytic delta-endotoxin, heat-stable enterotoxin A, Zot, accessory cholera enterotoxin, RTX protein, and thiol-activated cytolysin (Thomas and Ellar 1983; Huott et al. 1988; Trucksis et al. 1993; Fasano et al. 1995; Billington et al. 2000; Frey and Kuhnert 2002; Hood et al. 2010; Robb et al. 2016). Several of these viruses also encoded components of secretion systems (type II, type IVB Dot/Icm, and type VII), some of which may facilitate delivery of bacterial effectors into neighboring cells (Filloux 2022), although this is not an exclusive function of these secretion systems.

The top SCTLD indicator virus (1. SINT_DD_R2S43C_NODE_111), identified as a *Vibrio*-infecting *Caudoviricetes*, encoded a Tse2 homolog (Fig. 4). Tse2 is an NAD-dependent cytotoxin secreted via the Type VI secretion system (T6SS). To date, the natural secretion of Tse2 via T6SS has only been demonstrated in bacterial antagonism; however, intracellular expression experiments show that Tse2 can arrest growth and cause apoptosis-like or necrotic death to both prokaryotic and eukaryotic cells if delivered intercellularly (Hood et al. 2010). Notably, the coral pathogen *Vibrio coralliilyticus* pan-genome encodes two T6SSs, one mediating bacterial competition (T6SS1) and the other contributing more directly to targeting hosts through secretion of anti-eukaryotic effectors (T6SS2) (Mass et al. 2024). This suggests that Tse2 homologs could potentially contribute to direct coral cell lysis if delivered within coral tissues by *Vibrio* hosts. Similarly, toxin homologs in a virus predicted to infect *Flavobacteriia* (2. MCAV_DD_McD6_NODE_117) suggest a potential phage-mediated gene transfer mechanism to supply *Flavobacteriia* hosts with the capacity to compromise coral cell membranes and contribute to focal tissue necrosis. For example, the cytolytic delta-endotoxin (*cyt2Bb1*) is a pore-forming toxin that binds to eukaryotic membranes, leading to osmotic stress and cell lysis (Thomas and Ellar 1983). In contrast, heat-stable enterotoxin A (*YstA*) disrupts eukaryotic signaling by activating guanylate cyclase receptors, elevating cyclic GMP levels, and altering epithelial ion and fluid balance (Huott et al. 1988).

Filamentous viruses (*Inoviridae*) associated with SCTLD displayed the genomic potential to enhance bacterial invasion of coral tissues by compromising epithelial integrity. Notable virulence factors include *zot* and accessory cholera enterotoxin (*ace*), which disrupt epithelial barriers for pathogen invasion. Zot increases paracellular permeability by disrupting tight junctions between epithelial cells (Fasano et al. 1995), while Ace contributes to fluid secretion and epithelial barrier dysfunction (Trucksis et al. 1993).

RTX proteins are a diverse group of pore-forming toxins regulated by iron availability that promote adhesion, invasion, and cytotoxicity toward eukaryotic cells (Frey and Kuhnert 2002). Their presence in SCTLD-associated lysogenic conversion candidate viruses suggests that infected bacterial symbionts may acquire enhanced cytolytic capabilities, directly damaging coral cells or inhibiting competing microbes. Thiol-activated cytolysins, homologous to pneumolysin or seeligeriolysin, are cholesterol-dependent toxins that form large pores in eukaryotic membranes (Cockeran et al. 2002). These toxins can cause cell lysis and inflammation in animal hosts, and in corals, their activity could produce the cellular degradation and necrosis characteristic of SCTLD lesions.

Together, the discovery of these virulence-associated genes within SCTLD-associated bacteriophage genomes underscores the possibility that temperate and filamentous phages act as reservoirs of bacterial virulence, enhancing the cytotoxic potential of their hosts. Many of the virulence factors detected in SCTLD-associated phages, such as RTX proteins, Zot, and genes related to secretion or adhesion, are commonly carried on mobile genetic elements in pathogenic *Vibrio* strains, including prophages, genomic islands, and integrative conjugative elements (Wagner and Waldor 2002). In vibrios, this mobility drives rapid evolution of virulence and explains the substantial variability in pathogenic potential among strains (Faruque and Mekalanos 2003). Similarly, in our dataset, several of these virulence-associated functions were encoded exclusively on phage genomes from diseased coral samples, suggesting that lysogenic conversion may provide a mobile reservoir of toxins and fitness traits within coral-associated microbial communities. This parallel suggests that mechanisms well-characterized in known pathogens of other animals could also underlie the heterogeneous virulence and infection outcomes observed in SCTLD transmission experiments, linking phage-mediated gene transfer to the variability in coral susceptibility and disease progression.

### Conclusions and future directions

Collectively, our results highlight the potential role of lysogenic conversion in SCTLD pathogenesis. Temperate viruses enriched in diseased corals carry diverse putative virulence factors capable of disrupting eukaryotic cell membranes, interfering with tissue barriers, and modulating host-microbe interactions. This pattern suggests a conceptual model in which lysogenic conversion by temperate phages contributes to heterogeneity in disease outcomes. Depending on the presence or absence of specific prophages, their induction state, and horizontal gene transfer events, bacterial symbionts may gain or lose cytotoxic capabilities. In this framework, phages could underlie differential susceptibility within and among coral species, a spectrum of virulence and mortality (analogous to toxigenic vs. non-toxigenic *V. cholerae*) (Faruque and Mekalanos 2003). The specific viruses and virulence factors identified here provide concrete targets for experimental follow-up. For example, CRISPR-based editing of prophages, induction assays to trigger lytic cycles, or transcriptomic surveys to detect expression of these virulence genes could directly test the impact of phage-encoded functions on bacterial pathogenicity and tissue damage. Future work could also explore prophages and filamentous viruses of these SCTLD-associated bacterial hosts to identify genes, such as PftP4-like polypeptides, that selectively target their bacterial hosts when overexpressed (Weng et al. 2025). Such genes may provide a foundation for novel bacterial control strategies, offering potential tools for managing and detecting opportunistic pathogens in environmental settings.

## Declarations

### Data Availability

Original coral metagenomic sequence data for this project are available on NCBI (PRJNA1415870). The code used to conduct this analysis is publicly available on GitHub (https://github.com/Silveira-Lab/Wallace_SCTLD). R code and data are available on figshare (10.6084/m9.figshare.30853895 and 10.6084/m9.figshare.30853889).

### Author contributions

CBS, BAW, SMR, and LB conceptualized the study. EP and BU collected original samples. BAW and LB conducted sample processing and nucleic acid extractions. BAW conducted data retrieval, pipeline development, and analyses. BAW wrote the first draft, and all authors edited the manuscript.

## Acknowledgements

We would like to thank Sukanya Dayal, Yesmarie De La Flor, Dr. Valerie Paul, and the Smithsonian Marine Station staff, including Jay Houk, Katy Toth, and Woody Lee, for their support with coral collection. We also thank the researchers who have made their metagenomic datasets publicly available, as these resources were fundamental to the development of this project.

