## Supplemental Materials for "Temperate and filamentous bacteriophages as reservoirs of bacterial virulence in stony coral tissue loss disease"

### Supplementary Materials

### Supplementary Methods

#### *DNA Extractions*

For the original metagenomic data produced as part of this manuscript, three methods for DNA extractions were applied. For samples labeled “CS”, DNA extractions followed a previously described host depletion protocol (Wallace et al., 2024). Briefly, samples were crushed using a sterile mortar and pestle, suspended in 150 µL of sterile artificial seawater, and placed into a tube containing 0.2 g of 425- to 600-µm sterile glass beads (Sigma-Aldrich, St. Louis, MO). Samples were vortexed (VWR Analog Vortex Mixer; VWR, Radnor, PA) at speed 3 (~ 600 rpm) for 5 min to disrupt the larger coral and algal symbiont cells, and then treated with DNase I (20 U/ml final concentration) in DNAse I Buffer (Invitrogen, Waltham, MA) to degrade the coral host and algal DNA released in the previous step. DNAse activity was stopped with ethylenediaminetetraacetic acid (5 mM final concentration; Genesee Scientific, San Diego, CA), and the sample was filtered through an 8.0-µm membrane (Cytiva, Marlborough, MA) to allow passage of the bacteria and viruses. The flowthrough was concentrated with a 100 kDa Amicon Centrifugal Unit (Sigma-Aldrich, St. Louis, MO), and each side of the filter was rinsed and incubated at 56°C for 1 h (per side) in 200 µl Buffer T1 and 20 µl proteinase K from the NucleoSpin Tissue Kit (Macherey-Nagel Inc., Allentown, PA). DNA extraction proceeded from Step 3 of the kit, and DNA was eluted in 100 µl of PCR-grade water. A step-by-step description of the VBE protocol is publicly available on protocols.io (Varona et al., 2023).

DNA libraries for CB samples were prepared using the Nextera XT DNA Library Preparation Kit (Illumina) and IDT Unique Dual Indexes using 1 ng of DNA. Fragmentation was performed with Illumina Nextera XT fragmentation enzyme, followed by PCR amplification (12 cycles) and purification with AMpure magnetic Beads (Beckman Coulter). Libraries were quantified using a Qubit 4 fluorometer (ThermoFisher Scientific, Waltham, MA, USA) and a Qubit dsDNA HS Assay Kit (Invitrogen, Waltham, MA, USA). Libraries were sequenced on a NovaSeq platform 2 × 150 bp (Illumina, San Diego, CA, USA). Although all libraries generated sequencing data, sequencing depth varied substantially among samples, with 7 of 14 libraries yielding lower read counts that were accounted for in downstream abundance calculations.

For samples labeled ZB or MB, tools were cleaned with RNase Away (Thermo Fisher Scientific) and rinsed with Milli-Q water between samples. Coral tissue fragments (~1–2 cm) were placed into Zymo BashingBead tubes containing either 2 mm (ZB) or a mixture of 0.1 and 0.5 mm beads (MB), along with 800 µL of 1× phosphate-buffered saline (PBS). Samples were homogenized by vortexing the samples vertically for 45 min, after which homogenates were centrifuged at 5 × g for 30 s to pellet debris. A 200 µL aliquot of the supernatant was subsequently used for nucleic acid extraction using the HostZERO Microbial DNA Kit (Zymo Research, Irvine, CA) following the manufacturer's instructions.

All ZB and MB samples were submitted for sequencing at the University of Miami Miller School of Medicine Sequencing Core Hussman Institute for Human Genomics. DNA libraries for ZB and MB samples were prepared with 150-200ng of DNA (calculated with a Qubit). The NEBNext Ultra II DNA Library Prep Kit (Massachusetts, USA) was used with three PCR cycles. Libraries were sequenced on an Illumina Novaseq X Plus using a 10B-300 cycle kit to generate approximately 30 million paired-end 150 reads per sample.

### Supplementary Figures

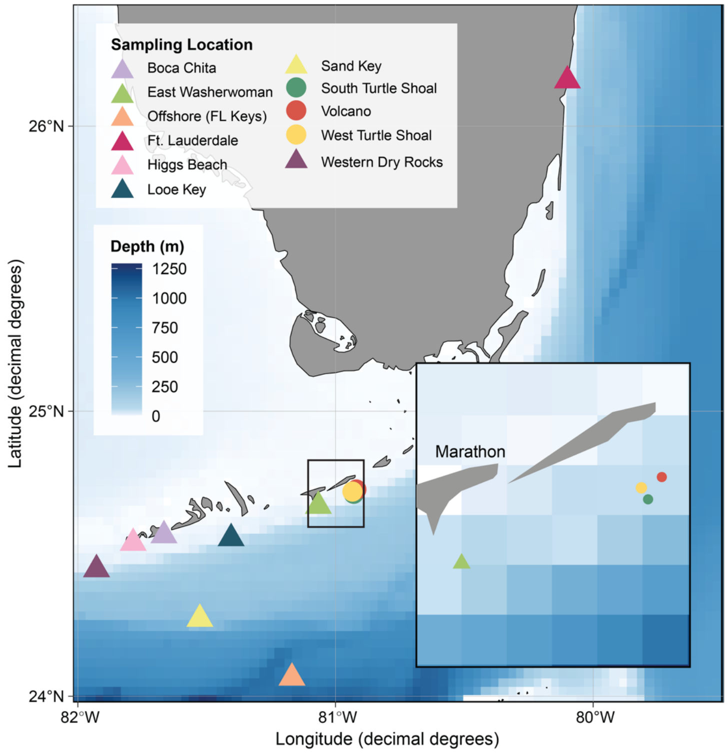


**Figure S1 | Sampling locations of coral metagenomes included in this study.** Locations where new coral samples were collected for this study are indicated on the map as circles. Additional metagenomic samples obtained from publicly available databases are indicated on the map as triangles. Only sites where GPS coordinates were provided by or could be inferred from the associated metadata are included. Thus, sites in the Kristin Jacobs Coral Reef Ecosystem Conservation Area (ECA) are not included. An inset of the map is provided to better show the locations of the original sampling sites for this project.

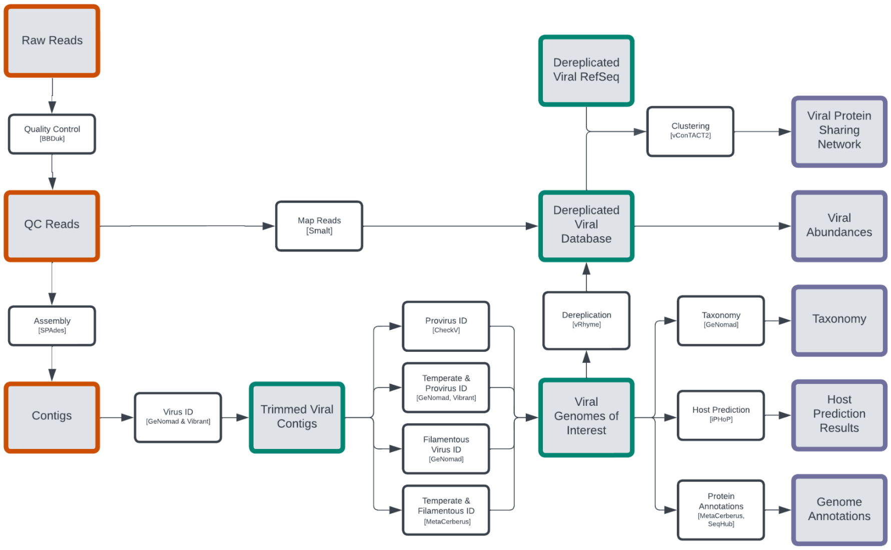


**Figure S2 | Bioinformatic workflow.** Bioinformatic workflow from QC and Assembly (orange) to viral contig identification and viral database curation (green) to viral annotation and host prediction (purple).

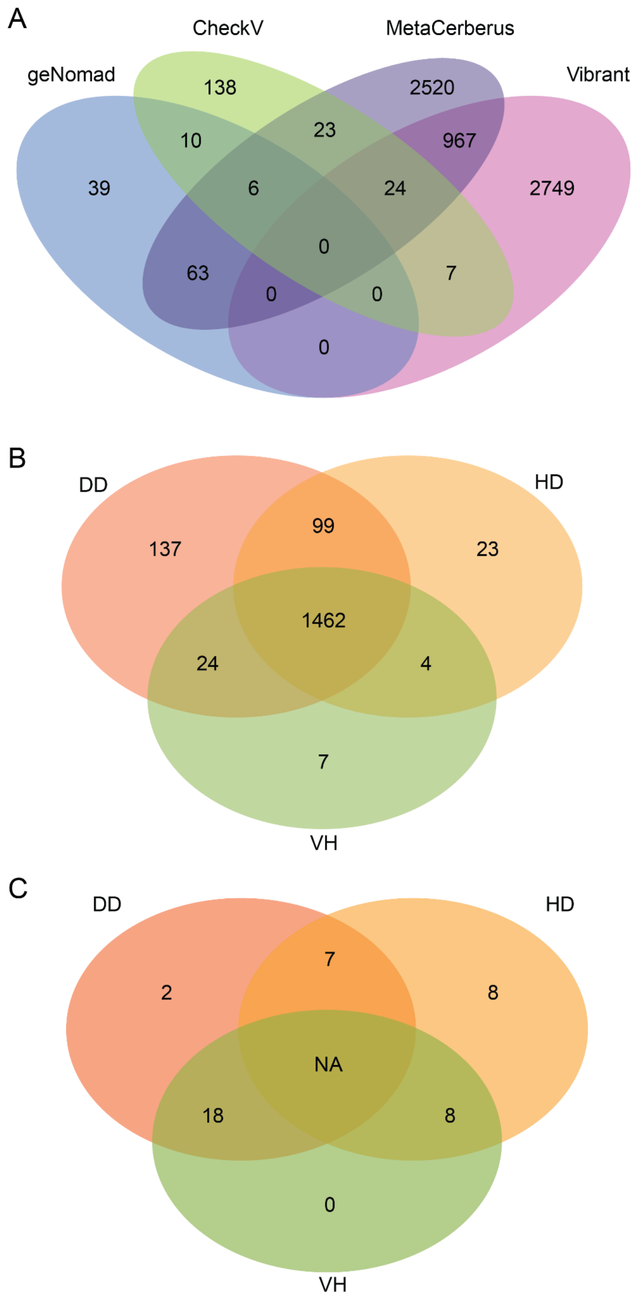


**Figure S3 | Venn diagrams of temperate virus counts.** (A) Counts of pre-dereplication temperate viruses identified by geNomad, CheckV, MetaCerberus, and Vibrant. (B) Counts of dereplicated temperate viruses present in DD, HD, and VH samples based on read-mapping. (C) Counts of temperate SCTLD indicator viruses of DD, HD, and VH groups.

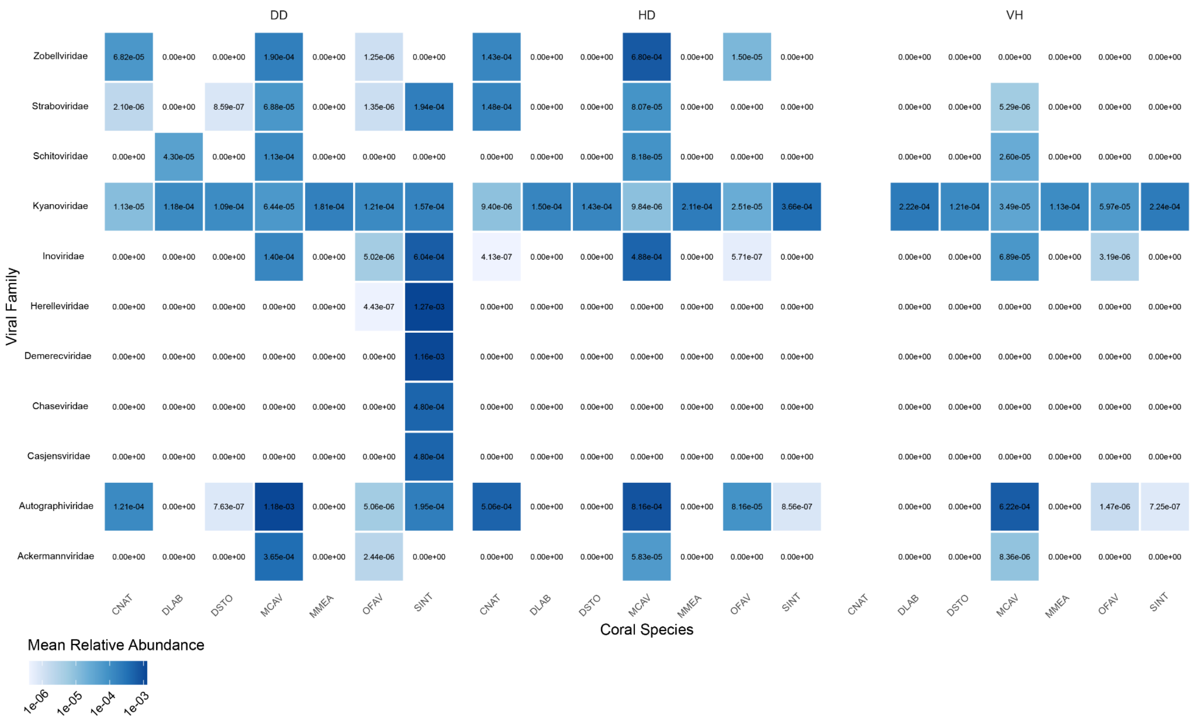


**Figure S4 | Lysogenic conversion candidate taxonomy by coral host species and disease state.** Relative fractional abundances of lysogenic conversion candidate families by disease state and coral host species. No healthy (VH) representatives of *Colpophyllia natans* (CNAT) were represented in this dataset.

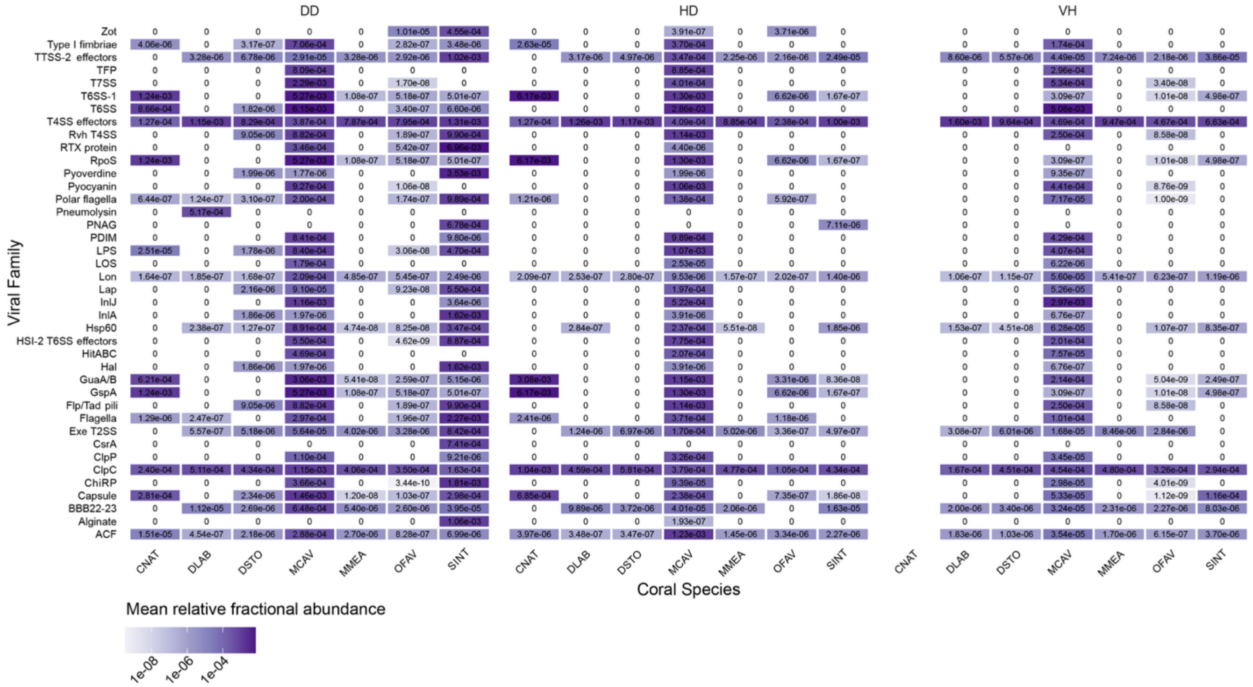


**Figure S5 | Virulence factors encoded by lysogenic conversion candidates across coral host species and disease state.** Relative fractional abundances of lysogenic conversion candidate-encoded virulence factors by disease state and coral host species. No healthy (VH) representatives of *Colpophyllia natans* (CNAT) were represented in this dataset.

### Supplementary Table Legends

**Table S1 | Metagenomic samples of SCTLD-infected and healthy corals.** Metadata for the coral-associated metagenomes used in this study. These metadata include the sequence run ID, sample ID, total bases, BioProject, BioSample, sample collection date, affiliation, consent, sample collection depth, country, geographic location, coral host species, reported health status, sample id number, sequencing instrument, a brief description of the DNA extraction method, latitude and longitude of sampling location (when available), library name, SRA study, and the original report.

**Table S2 | Significant indicator viruses.** Indicator species analysis was performed using the multipatt function in the R package indicspecies. This function uses a multi-level pattern analysis that calculates an indicator value for each group. Here we display the indicator viruses with an association statistic $\geq 0.5$. This revealed 43 viruses, 9 of which were significant ($p \leq 0.05$) indicators of diseased corals (DD samples only or the combination DD and HD samples).

**Table S3 | Predicted hosts of SCTLD-associated lysogenic conversion candidates in the network.** All viruses in the network (Virus_ID) are listed alongside their shortened name (Virus_ID2), the viral cluster they belong to (VC), their taxonomy (Tax) and the taxonomy method used (Tax_Method), their host assignment (Inferred_Host) and the method used to assign the host (Host_Method; from NCBI virus with known host, or a BLASTn search of host-flanking regions for proviruses), the consensus host of the viral cluster (VC_Host) with justification (VC_Host_Reason), and whether or not the virus was annotated as a provirus (Provirus).

**Table S4 | Sample distribution by coral species.** Coral host species and abbreviations are reported alongside the sample counts for each disease state.

**Table S5 | Viral family abundance and prevalence.** The mean abundance and prevalence of each viral family are reported for each disease state alongside the results of the Kruskal-Wallis rank-sum tests with Benjamini-Hochberg false discovery rate (FDR) adjustments and post hoc Dunn's tests with FDR correction. Significant values are bolded, and viral families with significant differences in abundance between disease states are highlighted in orange.

**Table S6 | Virulence gene abundance and prevalence.** The mean abundance and prevalence of each virulence factor are reported for each disease state alongside the results of the Kruskal-Wallis rank-sum tests with Benjamini-Hochberg false discovery rate (FDR) adjustments and post hoc Dunn's tests with FDR correction. Significant values are bolded, and viral families with significant differences in abundance between disease states are highlighted in orange.

**Table S7 | SCTLD-pathogenicity candidates.** Viruses linked to bacterial hosts previously implicated in SCTLD pathogenesis, members of viral families enriched in SCTLD-infected corals, and viruses encoding virulence genes enriched in SCTLD-infected corals are reported as potential SCTLD-pathogenicity candidates. For each virus, we report the viral cluster (VC), taxonomy (Tax), virus type (Type), and whether it encodes one or more virulence factors (VF). Virulence factors identified with Metacerberus are reported with the VF name (VFDB_Metacerberus_VF), VF identity (VFDB_Metacerberus_VF_ID), "best hit" accession (VFDB_Metacerberus_VF_Label), and description (VFDB_Metacerberus_VF_description).

Results from SeqHub are also included (SeqHub_VF_ID), along with the broader category of the SeqHub annotation (SeqHub_VF_Cat) and the host, where applicable.
